# Mechanisms for cross-neutralisation of diverse bat sarbecoviruses

**DOI:** 10.1101/2025.09.17.674891

**Authors:** Ayush Upadhyay, Jeffrey Seow, Yilmaz Alguel, Joseph Newman, Nazia Thakur, Abigail L. Hay, Jerry C.H. Tam, Andrea Nans, Richard J. Orton, Dalan Bailey, Peter Cherepanov, Katie J. Doores

## Abstract

The continuing evolution of SARS-CoV-2 variants of concern, and the increasing spillover potential of sarbecoviruses into the human population presents an important and urgent need to discover cross-reactive monoclonal antibodies (mAbs) for future therapeutic use and identify conserved neutralising epitopes that can be used for rationale design of broadly protective sarbecovirus vaccines. Here we study the neutralising epitopes on WIV-1 Spike of three mAbs that confer broad sarbecovirues and SARS-CoV-2 variant neutralisation, including XEC and JN.1. mAb V1WT_06 binds a highly conserved RBD site V epitope that is mediated by the heavy chain alone. V1WT_06 contact residues are highly conserved in circulating viruses suggesting that the epitope is evolutionarily and functionally constrained. mAbs V1WT_41 and VA14_26 bind overlapping RBD class 4 epitopes with differing angles of approach that impact on the degree of ACE2 competition. We show that neutralisation by these mAbs is maintained when virus entry is via Japanese horseshoe bat and Halcyon horseshoe bat ACE2. These mAbs are ideal candidates for therapeutic antibody development and inform the rational design of pan-coronavirus vaccines.

## Introduction

The virally encoded sarbecovirus surface glycoprotein, Spike, is critical for host cell attachment and entry, and is the major target for neutralising antibodies (nAbs) [1]. Spike is the predominant antigen target for both immunogen development and for antibody-based therapeutics against COVID-19. An increasing spillover potential of sarbecoviruses, and the continuing prevalence of emerging SARS-CoV-2 variants, presents an important and urgent need for identifying both cross-reactive monoclonal antibodies (mAbs) for therapeutic use and conserved neutralising epitopes that can inform rational design of broadly protective sarbecovirus vaccines. Since the beginning of the COVID-19 pandemic, the population has generated a high degree of humoral and cellular immunity to SARS-CoV-2, arising in response to both infection and vaccination [2–4]. An understanding of potential population-wide protection against new and emerging sarbecoviruses from animal hosts will be critical for future pandemic preparedness.

Sarbecoviruses can differ in their receptor usage for host cell entry. Whilst some viruses target a host-specific receptor for entry, others can utilise a range of receptors resulting in a wider host diversity [5–7]. Sarbecoviruses that use human angiotensin-converting enzyme 2 (ACE2) for entry are of most concern for zoonosis events as would likely require minimal adaptation. However, rapid virus adaptation [5] and the potential for expanded host range means that nAbs that also target animal viruses with zoonotic potential are of relevance and importance for future pandemic preparedness [5]. Furthermore, the continued selective pressure of humoral immunity is demonstrated by the on-going evolution of SARS-CoV-2 variants and the appearance of additional Spike mutations that reduce the potency of immune sera and escape therapeutic mAbs [8–12]. Therefore, there remains an urgent need to identify SARS-CoV-2 mAbs that can resist viral escape for development of improved and future-proofed antibody-based SARS-CoV-2 therapies.

The receptor binding domain (RBD) of Spike interacts with the ACE2 receptor on targets cells and is one of the dominant targets of nAbs generated in response to infection and vaccination [13–15]. Several epitopes on RBD have been identified. A dominant epitope is the highly variable receptor binding motif (RBM) where antibodies function by directly inhibiting the interaction of Spike with the ACE2 receptor (referred to Class 1 and Class 2 [14]). SARS-CoV-2 mutations frequently arise in this epitope due to the selective pressure of neutralising antibodies driving immune evasion [10, 16]. Other RBD epitopes distal to RBM include the CR3022 cryptic epitope (referred to as class 4 [14]), the N343-site (referred to as Class 3 [14]) and RBD site V [17].

We have shown that sarbecoviruses with a broad ACE2 tropism that includes human ACE2 (hu-ACE2) can be neutralised by SARS-CoV-2 convalescent and vaccinee sera [18–20]. In this study we also observed that some bat sarbecoviruses with more specialist ACE2 tropism were also weakly neutralised by these sera. Through screening a panel of neutralising monoclonal antibodies (mAbs) isolated from donors who were convalescent, vaccinated or experienced a breakthrough infection (BTI) [8, 21, 22], we identified several RBD-specific mAbs with broad binding and neutralising activity against bat sarbecoviruses belonging to multiple clades, including clades 1a, 1b, 3 and 5.

Here, we structurally characterised the RBD epitopes of three mAbs with the broadest SARS-CoV-2 variant neutralisation and sarbecovirus binding and studied their mechanisms of neutralisation in the context of ACE2 receptors from different hosts. We show that these mAbs target the RBD site V or class 4 epitopes and achieve their neutralisation breadth through binding to regions that are highly conserved between SARS-CoV-2 variants and bat sarbecoviruses. We identify a unique potently neutralising antibody that approaches the RBD site V with a binding mechanism and angle of approach that could potentially facilitate greater accessibility to the cryptic epitope. The binding epitopes of these mAbs make them ideal candidates for therapeutic antibody development and inform the rational design of pan-coronavirus vaccines.

## Results

### Cross-reactive mAbs bind non-competing epitopes on SARS-CoV-2 Spike

We identified three RBD-specific mAbs isolated from either a AZD1222 vaccinated individual (VA14_26) [22] or an individual, who was infected with the delta variant after BNT162b2 vaccination (V1WT_06 and V1WT_41) [8]. The three mAbs displayed broad cross-binding activity against SARS-CoV-2 variants and related bat sarbecoviruses including hu-ACE2 using viruses (BANAL-20-52, BANAL-20-236, WIV-1 and Rs4231), as well bat specific viruses that have more restricted bat ACE2 usage (Rc-o319, and RhGB07) (**Supplemental Figure 1A**) [20, 23, 24]. Fab versions of the three mAbs were cloned and expressed to facilitate detailed functional and structural characterisation of their neutralising epitopes. When comparing the binding of the Fab and IgGs against the RBDs of sarbecoviruses, there was a reduction in Fab Spike binding (**Figure 1A**) for V1WT_41, VA14_26 and V1WT_06 for some RBDs indicating that multivalency is important for binding to specific sarbecoviruses.

**Figure 1:**
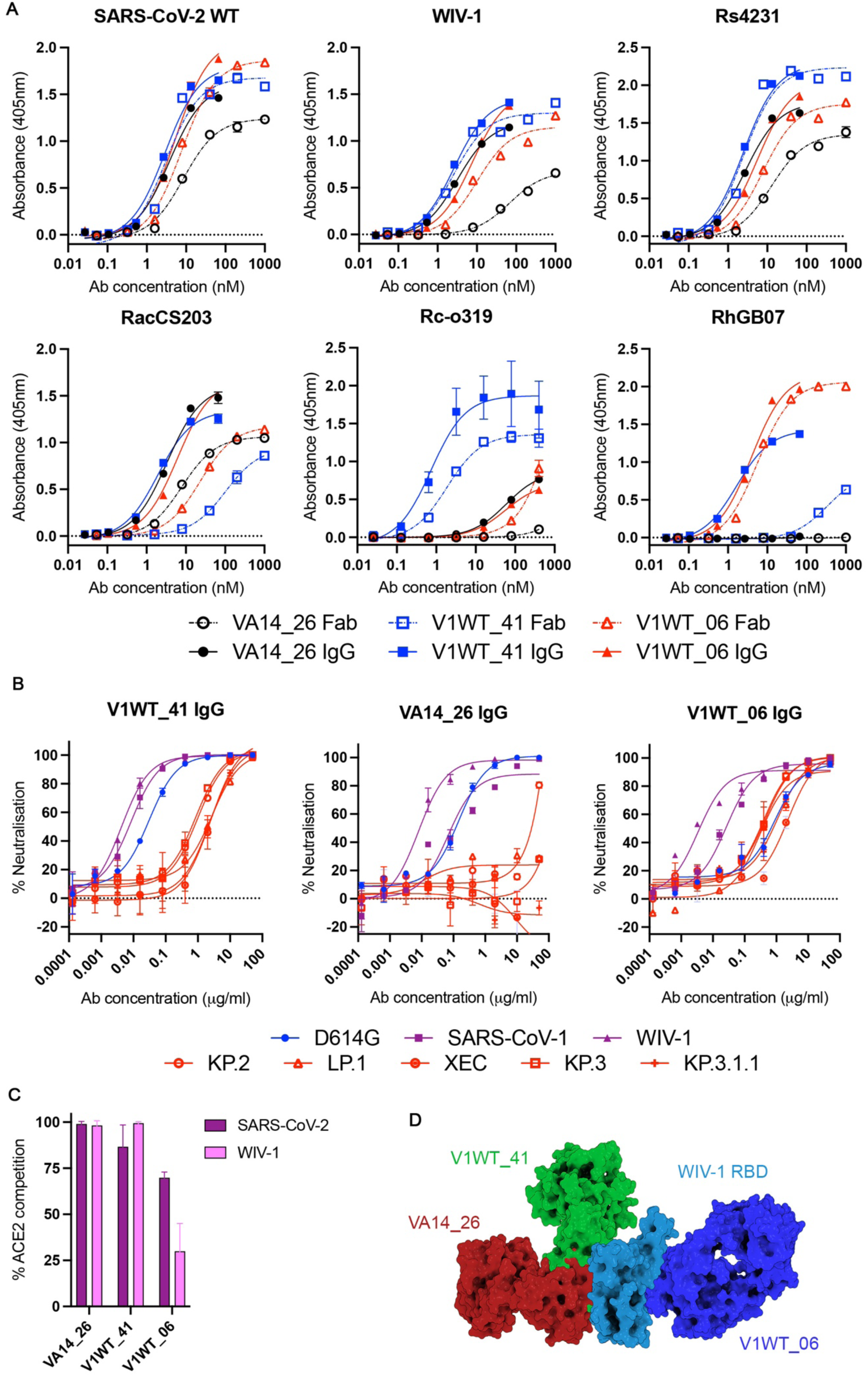
Binding and functional activity of V1WT_06, V1WT_41 and VA14_26. **A)** Binding of IgG and Fab to recombinant RBDs of Sarbecoviruses from different clades by ELISA including SARS-CoV-2 (clade 1b), WIV-1 (clade 1a), Rs4321 (clade 1a), RacCS203 (clade 1a), Rc-o319 (clade 5) and RhGB07 (clade 3). Experiments were run in duplicate and repeated twice. A representative assay is shown. Fab binding is shown with the dotted line and IgG binding with the full line. **B)** Neutralising activity of IgGs against SARS-CoV-2 and variants of concern as well as SARS-CoV-1 and WIV-1. **C)** Ability of IgGs to inhibit the interaction between cell surface human ACE2 and soluble SARS-CoV-2 and WIV-1 RBDs. IgGs were pre-incubated with fluorescently labelled RBD before addition to HeLa-ACE2 cells. The percentage reduction in mean fluorescence intensity is reported. Experiments were performed in duplicate. **D)** Superimposed 3D structures of the V1WT_06 (dark blue), V1WT_41 (green) and VA14_26 (red) around WIV-1 RBD (light blue, from PDB 8WLZ).

Neutralisation against the more recent SARS-CoV-2 variants was tested including KP.2, KP.3, LP.1, KP.3.1.1 and XEC and compared to the cross-neutralisation of SARS-CoV and WIV-1 (**Figure 1B**). V1WT_06 retained both breadth and potency against the VOCs. Whilst V1WT_41 retained breadth, there was between a 25- and 85-fold reduction in potency against the five SARS-CoV-2 variants tested. VA14_26 was most impacted by Spike mutations and was only able to weakly neutralise XEC. Interestingly, the three mAbs displayed higher neutralisation potency against the WIV-1 pseudotype compared to D614G and other SARS-CoV-2 variants.

To gain insight into the mechanisms of mAb cross-neutralisation, we performed an ACE2 competition assay using HeLa cells stably expressing hu-ACE2 and recombinant SARS-CoV-2 or WIV-1 RBDs (**Figure 1C**). V1WT_41 and VA14_26 competed with hu-ACE2 for binding to both SARS-CoV-2 and WIV-1 RBDs. In contrast, V1WT_06 showed only a partial hu-ACE2 competition with SARS-CoV-2 RBD and minimal competition with WIV-1 RBD (**Figure 1C**).

### Structural definition of mAb epitopes on WIV-1 Spike

To gain a detailed understanding of how the mAbs derive their broad cross-recognition, we imaged pre-fusion stabilised WIV-1 Spike ectodomain in the presence of the corresponding Fab fragments by cryo-EM. The WIV-1 Spike was stabilised in pre-fusion state by K969P and V970P mutations and fusion with the phage T4 fibritin trimerisation domain [25]. WIV-1 Spike was incubated with each Fab individually and the Spike:Fab complexes were applied onto the grids. Unbound WIV-1 Spike was observed to be predominantly in a ‘closed’ conformation where all RBDs where in the ‘down’ position (**Supplemental Figure 2**). A high degree of Spike protein disassembly was observed on the grids of all three Fab-Spike complexes and Fabs were only observed bound to the RBD within dissociated Spikes suggesting these mAbs may have destabilising effect on the trimer. The three mAbs bound distinct epitopes on the WIV-1 Spike, although some degree of overlap was observed in contact residues between VA14_26 and V1WT_41 (**Figure 1D**), which is consistent with the observed competition between VA14_26 and V1WT_41 binding for SARS-CoV-2 RBD (**Supplemental Figure S1B**).

### Characterisation of the V1WT_06 cryptic RBD epitope

V1WT_06, the broadest and most potent mAb studied, binds to a cryptic epitope overlapping with RBD site V that is distal to the ACE2 binding site and is only accessible when RBD is in the up conformation (**Figure 2A**). The location of the V1WT_06 epitope is consistent with the limited competition with hu-ACE2 (**Figure 1C**). V1WT_06 is only minimally mutated from germline, with the heavy and light chains containing 2.8% and 1.0% nucleotide mutation from germline, respectively. Compared to other reported SARS-CoV-2 mAbs (https://opig.stats.ox.ac.uk/webapps/covabdab/), V1WT_06 harbours an unusually long CDRH3. The 30-amino acid residue CDRH3 contains two cysteine residues that form a disulfide bond, three prolines and eight aromatic amino acids. The interaction between V1WT_06 Fab and WIV-1 RBD is mediated by the heavy chain alone, with the CDRH1, CDRH2 and CDRH3 all contributing to epitope recognition (**Figure 2B**). In particular, S49 in CDRH1 is in close proximity to RDB residue R462, the CDRH2 loop interacts in the region surrounding the E465 patch, and the long CDRH3 likely forms interactions with the residues on the antiparallel beta strands nestling within a shallow pocket on the RBD (**Supplemental Figure 3A**).

**Figure 2:**
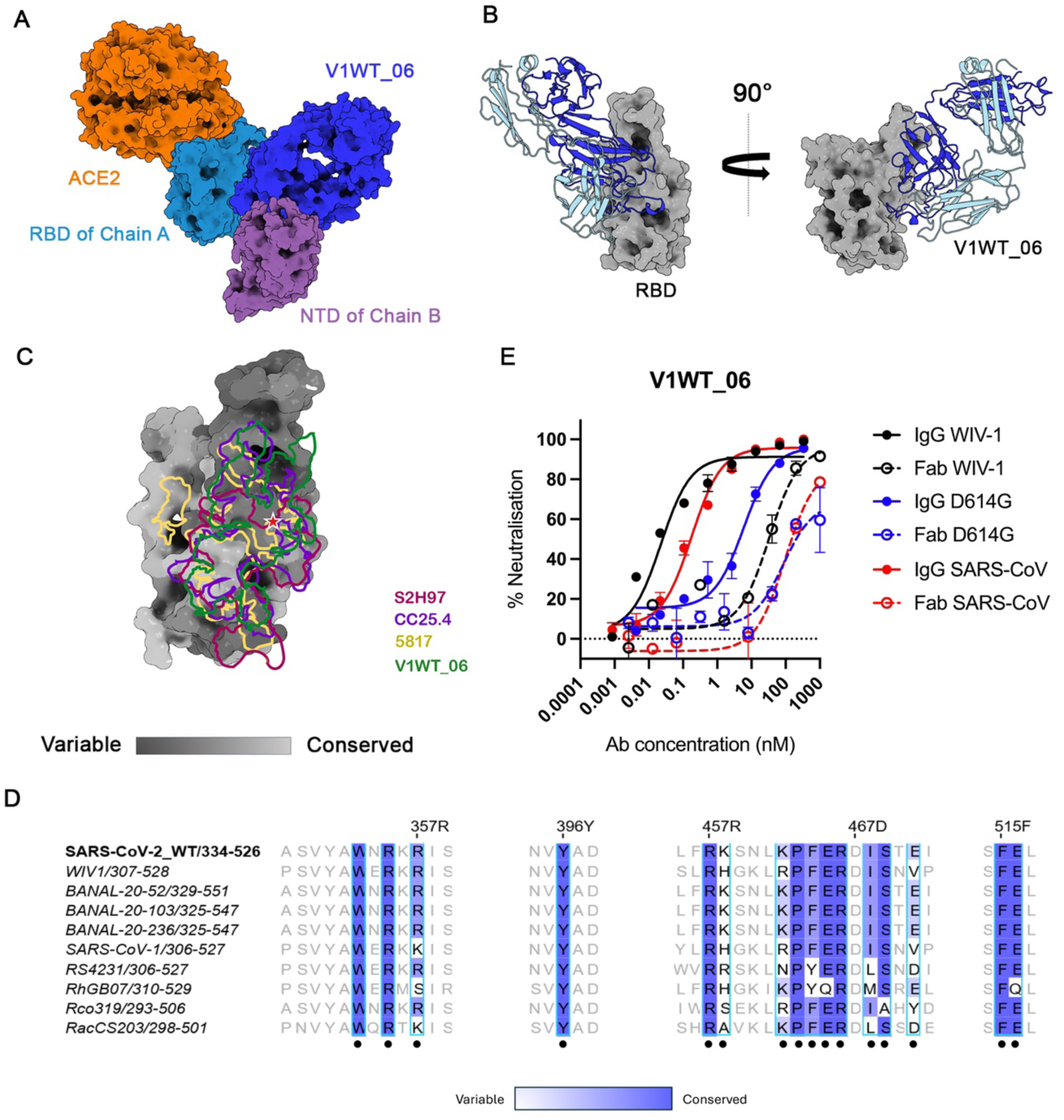
V1WT_06 binds to the RBD site V epitope using the heavy chain alone. **A)** A closeup view of the superimposed structure of V1WT_06 bound to WIV-1 RBD on a trimeric spike (from PDB: 8WLZ). A clash with NTD of chain B is observed when V1WT_06 Fab-RBD complex is aligned onto RBD in up conformation (chain A) in trimeric WIV-1 spike protein. Notably, the binding site is inaccessible in the RBD-Down conformation. **B)** V1WT_06 heavy chain only interaction with WIV-1 RBD (grey). Heavy and light chains are shown with the dark and light blue ribbons, respectively. **C)** V1WT_06 epitope footprint overlayed with epitope footprints of other site V neutralising antibodies including S2H97, CC25.4, and 5817 [17, 27, 28]. SARS-CoV-2 RBD (from PDB: 8m0j) is coloured based on the amino acid conservation across 87 bat sarbecoviruses [6, 23, 75, 76]. **D)** Sequence alignment of a panel of bat sarbecovirus RBDs. Contact residues of V1WT_06 mAb on the WIV-1-RBD are indicated with black dots. The colouring scheme describes the percent identity (PID) between the sequences as implemented in Jalview 2.11.4.0. **E)** Comparison in neutralisation potency of V1WT_06 Fab and IgG against SARS-CoV-2 (D614G), SARS-CoV-1 and WIV-1.

Comparison of the V1WT_06 epitope with other RBD site V SARS-CoV-2 mAbs shows a similar epitope footprint to mAbs S2H97 [17], WRAIR-2063 [26], 5817 [27] and CC.25.4 [28] (**Figure 2C**). The epitope of RBD site V mAbs is centred around the “E465 patch” which has been reported to be mutationally constrained and evolutionarily conserved [29]. S2H97, 5817 and WRAIR-2063 have shorter CDRH3 regions (11, 18 and 17 amino acids, respectively) than V1WT_06, and their light chains also engage in epitope recognition [17, 26, 27]. These observations indicate that extended CDRH3 is not a specific antibody requirement for targeting of this RBD epitope, a feature observed for some HIV-1 bnAbs [30–32].

The unchanged potency of V1WT_06 against KP.2, KP.3, KP.3.1.1, LP.1 and XEC compared to D614G is due to the conservation of contact residues within the V1WT_06 epitope (**Supplemental Figure 3B**). Analysis of the GISAID database shows that mutations at V1WT_06 contact sites on RBD are rarely seen in circulating Spikes (**Supplemental Figure 4**). Indeed, mutations in the RBD site V epitope have been shown to severely impact protein expression and might suggest that there are limited escape pathways available [17]. Alignment of WIV-1 contact residues with other bat sarbecovirus Spikes show that despite broad reactivity, not all WIV-1 RBD contact residues are conserved including R357, R462, F464, E465, I468, V471 (**Figure 2D**) suggesting an ability to tolerate diversity within the epitope footprint which may contribute to neutralisation breadth. Furthermore, two WIV-1 contact residues differ compared to SARS-CoV-2 RBD, R462K and V471E, which may explain the reduced neutralisation potency against SARS-CoV-2 compared to WIV-1 (**Figure 1B**).

Interestingly, for other Class V nAbs the RBD cross-binding activity did not always equate to neutralisation. For example, WRAIR-2063 had no neutralising activity against WIV-1 and SHC014 despite good binding to their RBDs [17]. Additionally, S2H97, although could neutralise WIV-1, had a lower potency (∼12 μg/mL) compared to SARS-CoV-2 (∼3 μg/mL) [26]. Despite the considerable overlap in contact residues between site V mAbs (**Figure 2C**), V1WT_06 and S2H97 bind RBD with differing angles of approach (**Supplemental Figure 3C**). The majority of site V mAbs bind the epitope perpendicular to RBD when modelled onto Spike [26]. However, the heavy chain mediated binding of V1WT_06 alters the Fab approach to be more upright in relation to RBD when in the “up” conformation (**Figure 2A**). This altered binding angle and heavy chain interaction would likely make the site V epitope more accessible to V1WT_06 compared to other site V antibodies which may enhance neutralisation potency.

The differences in neutralisation potency may also relate to distinct mechanisms of neutralisation. S2H97 was shown to cause rapid and premature refolding of SARS-CoV-2 Spike to the post-fusion conformation and S1 shedding [17]. However, WIV-1 and SARS-CoV lack the Spike cleavage site, which may explain the minimal neutralisation potency of S2H97 against these viruses. To determine whether V1WT_06 might share a similar neutralisation mechanism to S2H97, we measured the ability of mAbs to facilitate shedding S1 from HEK 293T cells expressing SARS-CoV-2 Spike by flow cytometry [17]. Positive control mAb (P008_067) showed a reduction in binding indicative of S1 shedding from the cell surface, whereas V1WT_06 showed no evidence of causing shedding of S1 (**Supplemental Figure 5**). By contrast to P008_067, V1WT_06 showed no evidence of S1 shedding in this assay. Modelling V1WT_06 bound to SARS-CoV-2 Spike with RBD in the “up” conformation predicts a steric clash between the Fab and the NTD from neighbouring S1 chain (**Figure 2A**), similar to that observed for CC.25.4 [28] and 6D6 [33]. Wedging of the Fab between S1 subunits, would conformationally restrict the Spike and/or lead to trimer destabilisation, as observed in our cryo-EM analysis. Indeed, the bulkier IgG showed a 1275- and 45-fold more potent neutralisation of WIV-1 and D614G, respectively, than the Fab alone (**Figure 2E**) despite there being only a 6.3- and 1.3-fold increase in binding by ELISA.

Overall, V1WT_06 binds an RBD site V epitope with a novel heavy chain only mode of binding that allows for antibody approach from a more favourable binding angle, leading to high potency and breadth.

### VA14_26 and V1WT_41 bind epitopes overlapping with RBD Class 4 mAbs

The epitopes of VA14_26 and V1WT_41 partially overlap each other and are adjacent to the ACE2 binding site (**Figures 3A-C**). Their epitopes also overlap with that of CR3022 [34] and other RBD class 4 mAbs [14] including 10-40 [35] and ADI-62113 [36] (**Figure 3B**). VA14_26 Fab binds RBD using both the heavy and light chains with 1427 Å^2^ buried surface area (**Supplemental Figure 6B & 6C**). Although the cryo-EM structure does not predict a direct clash, when superimposed to the RBD, the Fab and ACE2 would come in immediate contact to achieve a backbone separation of <10 Å at several positions (**Figure 3B**). Thus, a direct interaction and/or allostery may help explain the observed competition between the Fab and receptor (**Figure 1C**). Notably, the VA14_26 Fab construct displayed 109-fold reduced neutralisation potency compared to the full IgG (**Supplemental Figure 6A**). Thus, the mechanism of neutralisation by this bnAb may involve bivalent binding to the Spike, which could more effectively occlude the RBD from the receptor.

**Figure 3.**
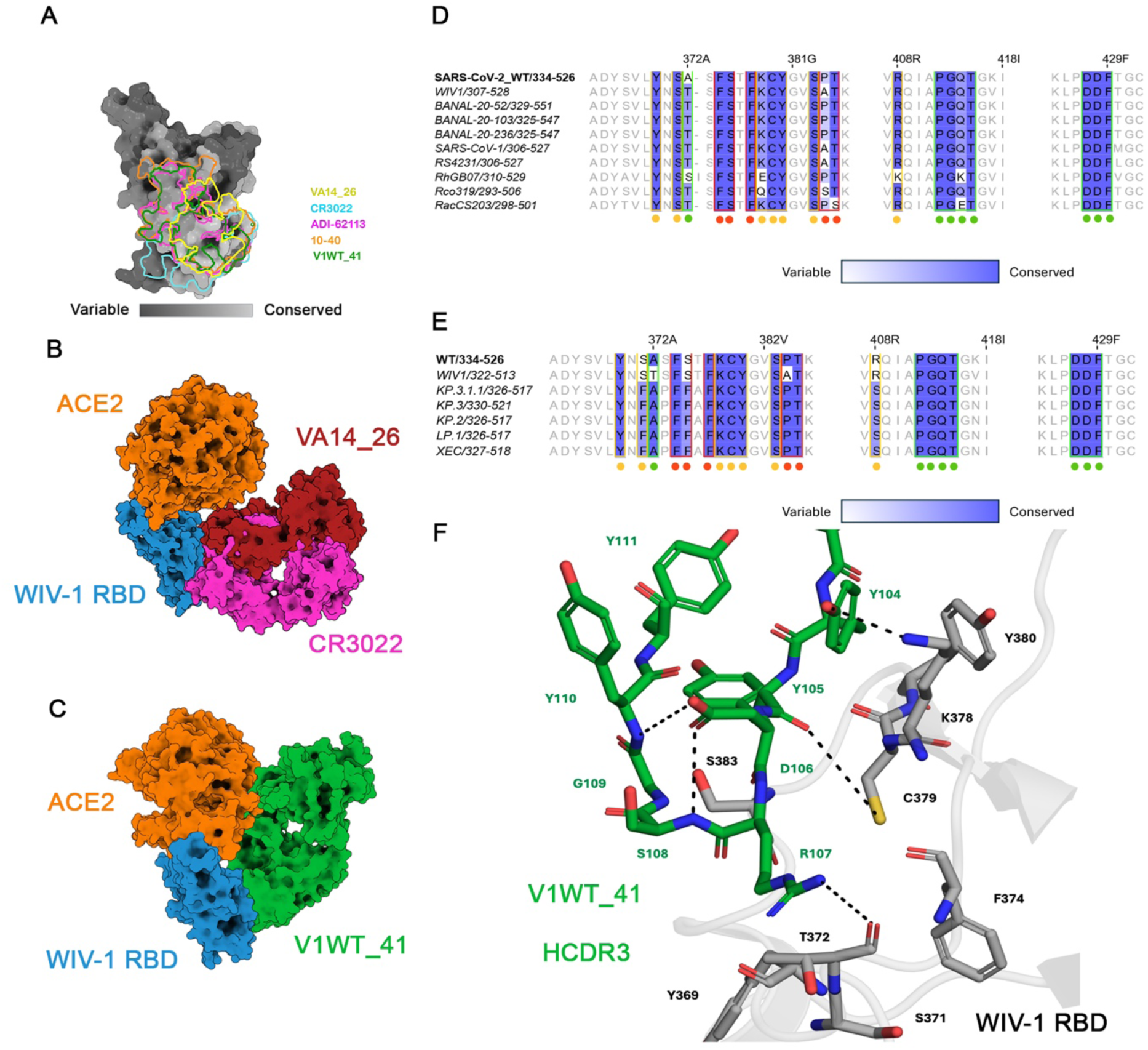
VA14_26 and V1WT_41 bind epitopes that overlap with the RBD class 4 epitope. **A)** VA14_26 and V1WT_41 antibody footprints overlayed with the epitope footprints of other RBD class 4 neutralising antibodies including CR3022, 10-40, and ADI-62113 [34] [35] [36]. SARS-CoV-2 RBD (from PDB: 8m0j) is coloured based on the amino acid conservation across 87 bat sarbecoviruses [6, 23, 75, 76]. **B)** Superimposed 3D structure depicting the binding of VA14_26 (Red) to WIV-1 RBD (blue) in relation to ACE2 (orange, from PDB: 8WLZ) and CR3022 (pink, from PDB: 6W41). **C)** Superimposed 3D structure depicting the binding of V1WT_41 (green) to WIV-1 RBD (blue) in relation to ACE2 (orange, from PDB: 8WLZ). **D)** Sequence alignment of a panel of bat sarbecovirus RBDs. Contact residues for VA14_26 and V1WT_41 are indicated by red and green dots and outline, respectively. Shared contact residues are shown with yellow dots and outline. **E)** Sequence alignment of a panel of SARS-CoV-2 variant RBDs. Contact residues for VA14_26 and V1WT_41 are indicated by red and green dots and outline, respectively. Shared contact residues are shown with yellow dots and outline. The colouring scheme describes the percent identity (PID) between the sequences as implemented in Jalview 2.11.4.0. **F)** A close-up view of YYDRSG motif in V1WT_41 HCDR3 (green) in contact with WIV-1 RBD (grey), shown in stick representation, renumbered based on alignment to SARS-CoV-2 [35, 36]. Presence of G109 breaks the H-bonding pattern in the beta hairpin to create a G1 β-bulge. This further enables D106 side chain to make hydrogen bond with the amides in the protein backbone, thereby stabilising the local structure. Combined this accentuates a right-handed twist found in G1 β-bulge [77]. Twisting of the β-hairpin is thought to contribute towards stabilisation of RBD-Fab interaction in several other public mAbs with YYDRxG motif [35, 36].

Of the three mAbs studied here, VA14_26 has the narrowest cross-reactivity with both SARS-CoV-2 variants (**Figure 1B**) and sarbecoviruses (**Figure 1A**), which is consistent with differences at critical RBD contact positions (**Figure 3D and 3E**). Thus, the lack of VA14_26 neutralisation against variants KP.2, KP.3, KP.3.1.1, LP.1 and XEC is likely due to the appearance of the Spike mutations S371F, S375F and R408S which abolish interactions with residues in the CDRH1 and CDRH3 (**Supplemental Figure 6B & 6C**). Whereas lack of binding to RhGB07 is likely due to a difference in amino acids at R378E and R408K (**Figure 3D**).

The epitope footprint of Fab V1WT_41 overlaps with VA14_26 but is larger in size (buried surface area of 1795 Å^2^), extending towards the RBM and incorporating additional WIV-1 residues at P412, G413, Q414, T415, D427 and D428, F429 (**Figure 3C and 3D**). Although the V1WT_41 footprint does not overlap directly with the ACE2 footprint, the more downward angle of Fab binding relative to Spike when embedded upright in a membrane means there is a significant steric clash with hu-ACE2 consistent with the ACE2 competition data (**Figure 3C and 1C**).

The CDRH3 of V1WT_41 is 22 amino acids long and contains a YYDRxG motif that has previously been shown to be important for a subset of mAbs that recognise the Class 4 RBD site, including 10-40 [35], COVA1-16 [37] and ADI-62113 [36]. The YYDRxG motif is encoded by the D3-22 allele, an allele that is frequently used in the human antibody repertoire [35, 36]. Structural analysis of ADI-62113 shows the YYDRxG motif forms a conserved structure at the tip of the CDRH3 loop that interacts with RBD and is stabilised by a preceding β-bulge [36]. The V1WT_41-RBD interface involves CDRH3 residues Y104, D106, R107 and G109 with RBD residues K378, C379, S371 and S383 (**Figure 3F**). The CDRH3 and framework-2 regions also interact with R408 (**Supplemental Figures 6D & 6E)**.

V1WT_41 has a broader VOC and sarbecovirus cross-reactivity than VA14_26. Although there is variation at WIV-1 contact sites for the other sarbecoviruses studied here, cross-reactivity is maintained (**Figure 1A and Figures 3D & 3E**). Spike mutations S371F and R408S present in SARS-CoV-2 variants KP.2, KP.3, KP.3.1.1, LP.1 and XEC reduce neutralisation potency (25-80-fold increase in IC_50_) but do not abolish neutralisation activity entirely (**Figure 1B**). The larger footprint of V1WT_41 compared to VA14_26 may allow more sequence variation to be tolerated resulting in broader cross-reactivity. In general, the V1WT_41 contact residues are very well conserved in the sarbecovirus genus (**Figure 3D**) suggesting that this region may have strong functional constraints that prevent mutation.

Higher resolution structural analysis of Fab 10-40 (another YYDRxG mAb) with both WIV-1 and SARS-CoV-2 RBDs revealed a higher number of Fab contacts with WIV-1 compared to SARS-CoV-2 leading to higher neutralisation potency against WIV-1 [35]. Similarly, the higher neutralisation potency of V1WT-41 against WIV-1 may be related to a difference in contact residue T372A in SARS-CoV-2 (**Figure 1B**). Despite the disassembly of soluble WIV-1 Spike ectodomain observed for V1WT_41 and VA14_26 by cryo-EM, neither mAbs were able to facilitate shedding of the S1 from SARS-CoV-2 Spike expressed on the surface of 293 T cells (**Supplemental Figure 5**).

Overall, VA14_26 and V1WT_41 bind a conserved and overlapping epitope with differing angles of approach that alters steric clash with hu-ACE2.

### Functional activity of mAbs when host cell entry is via Japanese horseshoe and Halcyon horseshoe bat ACE2

Previous studies that identified cross-reactive mAbs have generally focussed on neutralisation of sarbecoviruses entering target cells via hu-ACE2. As not all the sarbecoviruses tested are able to use hu-ACE2 for cell entry [23, 24], we compared the ability of mAbs to both compete with different species ACE2s for RBD binding and to neutralise viruses when entry was via a non-human ACE2, including Japanese horseshoe bat ACE2 (*Rhinolophus cornutus, Rh. cornutus*-ACE2) and Halcyon horseshoe bat ACE2 (*Rhinolophus alcyone, Rh. alcyone*-ACE2) (**Figure 4**).

**Figure 4:**
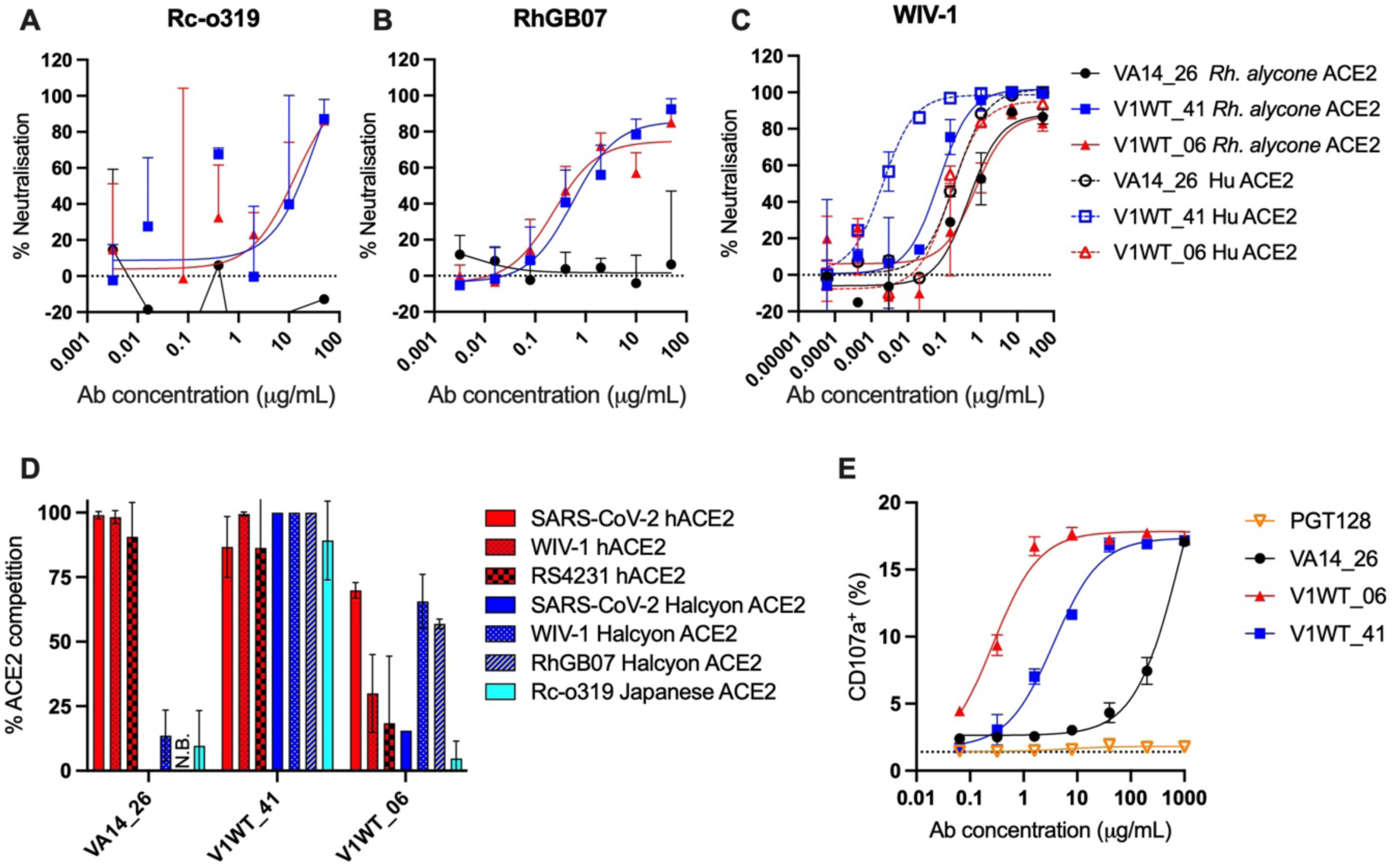
Mechanism and functional activity of V1WT_06, V1WT_41 and VA14_26. Neutralisation of sarbecoviruses through entry via different host ACE2s. Neutralisation of **A)** Rc-o319 and **B)** RhGB07 were measured against entry via *Rh. cornutus* ACE2 and *Rh. alcyone* ACE2, respectively, which were transiently expressed in HEK293T/17 cells. **C)** Neutralisation of WIV-1 was compared between entry via hu-ACE2 (dotted line and open symbols) and *Rh. alcyone* ACE2 (full line and closed symbols). Neutralisation was measured using HEK 293T cells transiently expressing either *Human* ACE2 or *Rh. alcyone* ACE2. **D)** Ability of IgGs to inhibit the interaction between cell surface ACE2 (including hu-ACE2, *Rh. cornutus* ACE2 and *Rh. alcyone* ACE2) and soluble RBD (including SARS-CoV-2, WIV-1, Rs4231, RhBG07 and Rc-o319 RBDs). IgGs were pre-incubated with fluorescently labelled RBD before addition to HeLa-ACE2 cells. N.B=No binding of IgG to the RBD. The percentage reduction in mean fluorescence intensity is reported. Experiments were performed in duplicate. **E)** Antibody dependant cellular cytotoxicity (ADCC) of IgGs measured through the surface expression of CD107a on activated NK cells (NK-92 cells expressing CD16) [38]. HIV-1 specific neutralising antibody, PGT128 is used as a negative control [31].

First, we measured the ability of the three mAbs to neutralise RhGB07 and Rc-o319 entering via *Rh. alcyone* ACE2 and *Rh. cornutus* ACE2, respectively. Consistent with the cross-binding reactivity of mAbs (**Figure 1A**), V1WT_06 and V1WT_41 were able to neutralise both RhGB07 and Rc-o319, whilst VA14_26 showed no neutralisation (**Figure 4A & 4B**). Next, we compared neutralisation of WIV-1 when cell entry was mediated by *Rh. alcyone* ACE2 or hu-ACE2. The neutralisation of WIV-1 by V1WT_06 and VA14_26 appeared independent of the host origin of the ACE2 receptor used whereas V1WT_41 was more potent at neutralising WIV-1 when entry was mediated by hu-ACE2 (**Figure 4C**).

To complement the neutralisation studies, we compared the ability of mAbs to inhibit binding of SARS-CoV-2, WIV-1, Rc-o319 RBDs to *Rh. alcyone* ACE2 and binding of RhGB07 RBD to *Rh. cornutus* ACE2. As expected, there was minimal ACE2 competition for site V binding mAb V1WT_06. V1WT_41 competed with ACE2 binding for all RBD and ACE2 pairings (**Figure 4D**) confirming that inhibition of ACE2 binding is the likely mechanism of V1WT_41 neutralisation. However, the more potent V1WT_41 neutralisation of WIV-1 entering through hu-ACE2 does not appear to be due to more efficient ACE2 competition. In contrast, despite VA14_26 showing potent neutralisation of WIV-1 entry via both hu-ACE2 and *Rh. alycone*-ACE2, VA14_26 was unable to compete with *Rh. alycone*-ACE2 binding for WIV-1 and SARS-CoV-2 (**Figure 4D**) suggesting the mechanism of VA14_26 neutralisation is only partially dependent on ACE2 competition. Overall, these data show that neutralisation breadth is conserved for different sarbecovirus and host ACE2 combinations.

### mAbs show potent ADCC activity

Finally, we determined whether these mAbs could facilitate antibody-dependent cellular cytotoxicity (ADCC) against ancestral SARS-CoV-2 [38, 39]. All three cross-reactive mAbs demonstrated Wuhan-1 specific potent degranulation of an NK cell line expressing CD16 (NK-92) detected through measurement of CD107a expression by flow cytometry (**Figure 4E**) whereas the negative control, HIV-specific mAb PGT128, showed no activity. VA14_26 had the lowest activity which is consistent with the lower binding affinity to Spike. Interestingly, mAbs 5817 and S2H97, which bind an epitope over lapping with V1WT_06 were reported not to mediate potent ADCC activity [17, 40] suggesting the angle of antibody binding can also play a role in effector function activity. Overall, in a scenario where cross-sarbecovirus binding exists in the absence of neutralisation, effector functions of mAbs generated following SARS-CoV-2 infection/vaccination could still provide some level of protection upon viral exposure [39].

## Discussion

Here we describe the neutralising epitopes of three novel mAbs that have broad SARS-CoV-2 variant and sarbecovirus cross-binding and cross-neutralisation. The continued evolution of SARS-CoV-2 Spike and the sustained threat of novel sarbecoviruses zoonosis into the human population warrants a more complete understanding of neutralising epitopes that are highly conserved across sarbecoviruses. Identification of broad antibodies and conserved epitopes will have application to development of next generation antibody-based therapeutics and for rational vaccine design.

mAbs with a high degree of sarbecovirus cross-reactivity appear to be relatively rare within the immune repertoire of SARS-CoV-2 infected and/or vaccinated individuals. However, they are present in donors with minimal and differing SARS-CoV-2 exposure histories. Only two AZD1222 vaccinations were required to elicit VA14_26 [22], whilst V1WT_41 and V1WT_06 were isolated from a donor that received two BNT162b2 vaccinations and experienced a breakthrough infection with delta VOC [8]. Other studies reporting cross-reactive mAbs to RBD class 4 and site V were also isolated from individuals with minimal SARS-CoV-2 exposures [13, 14, 17, 41]. This observation differs from HIV-1 infection where chronic antigen stimulation and extensive somatic hypermutation is required to achieve neutralisation breadth against multiple HIV-1 strains [30, 42].

While the RBM is highly variable between sarbecoviruses and SARS-CoV-2 variants, the RBD does harbor conserved epitopes. RBD site V is highly conserved across sarbecoviruses, and deep mutational scanning has shown that mutation around the E465 patch severely impacts on RBD expression [29], highlighting the evolutionary and functional constraints on the site V epitope. V1WT_06 is unique compared to other site V nAbs as binding is mediated by the heavy chain alone. This property enables the mAb to approach the cryptic site at an oblique angle compared to other site V nAbs, potentially permitting access to a transitional state of Spike with relatively narrow opening between the NTD and RBD on neighbouring protomers. HIV-1 broadly neutralising antibodies (bnAbs) that target the V2-apex bnAb epitope are also dominated by heavy chain interactions where a long CDRH3 penetrates through the glycan shield to contact conserved residues at the base of the glycans [30, 43]. However, the light chain also makes minor contacts with HIV-1 Env [32] but there is flexibility in terms of light chain pairing [44]. It may be possible to adapt vaccination strategies engineered to selectively elicit HIV-1 antibodies with long CDRH3s to engineer immunogens that elicit V1WT_06-like nAbs against the RBD site V epitope [45–47].

The mechanism of virus neutralisation by mAbs to site V is not fully understood. Whilst mAbs such as S2H97 are able to facilitate S1 shedding [17], not all sarbecoviruses Spikes encode an S1/S2 furin cleavage site, which is conceivably required for physical loss of the S1 subunit [48]. Our data suggests that binding of non-neutralising antibodies to the site V epitope may still facilitate effector functions such as ADCC which are important for protection from infection [39]. V1WT_06 was able to neutralise all sarbeoviruses tested regardless of the S1/S2 furin cleavage site indicating an alternative neutralising mechanism to S1 shedding. Modelling of the V1WT_06/RBD bound to trimeric Spike suggests Fab binding drives a wedge between RBD and NTD on neighbouring protomers that may destabilise the Spike protein leading to the Spike disassembly observed in our cryo-EM data.

The remarkable conservation of the RBD class 4 site is thought to be due to its role in stabilising the RBD-RBD interface when RBD is in the down conformation and it can also form contacts to the S2 domain [34]. The limited exposure of this epitope and the subsequent low frequency of mAbs targeting this domain might also reduce the immune selective pressure on this epitope leading to limited variation in this region. V1WT_41 belongs to the well-described public antibody class encoding a YYDRxG motif in the CDRH3 region [13, 35, 36]. The mechanism of V1WT_41 neutralisation is through sterically restricting ACE2 binding, a mechanism we show is conserved when sarbecovirus entry is mediated through ACE2 of Japanese (*Rh. cornutus*) and Halcyon (*Rh. alycone*) horseshoe bats.

The VA14_26 epitope overlaps with the CR3022 epitope but is considerably broader in neutralisation than CR3022 [22, 34]. Broadly reactive mAbs that target the CR3022 epitope are rare. Examples include C118 which achieves cross-neutralisation through a long CDRH3 reaching the conserved patch on RBD as well as contacting backbone residues in RBD allowing tolerance of some degree of variation [49]. The mechanism of VA14_26 neutralisation is less clear than V1WT_41 as the more perpendicular angle of Fab binding and the epitope position suggests that the steric clash with hu-ACE2 is minimal. Furthermore, VA14_26 shows limited competition with *Rh. alycone*-ACE2 for WIV-1 RBD binding compared to human-ACE2 but showed similar neutralisation potency when cell entry mediated by *Rh. alycone*-ACE2 or human-ACE2. As VA14_26 showed no evidence of causing S1 shedding, the mechanism of neutralisation is likely through Spike destabilisation.

The low frequency of mAbs to RBD class 4 and site V is likely explained transient exposure of their epitopes only when RBD is in the “up” conformation. By contrast, the immunodominant neutralising epitopes on RBD map to the RBM (including Class 1 and 2 epitopes) [13, 14, 17]. Antibodies targeting these epitopes tend to be highly potent, but their epitopes are more highly mutated, particularly in SARS-CoV-2 variants, due to the selective pressure of nAbs leading to high sequence variation and minimal breadth. The key challenge for rational immunogen design is to engineer immunogens that selectively expose or unmask these subdominant yet highly conserved class 4 and site V RBD epitopes. When considering these epitopes in the context of sarbecoviruses more broadly, the WIV-1 Spike is mostly found with RBD in the “down” conformation [50] suggesting immunogens based on the full Spike protein will be poor at inducing such antibodies. Several groups have investigated the use of mosaic RBD nanoparticle vaccines for elicitation of broadly reactive RBD-binding antibodies in mice. The multivalent presentation of mixed RBDs was shown to preferentially generate antibodies towards the conserved Class 1/4 [51] and site V epitopes [52]. In particular, a self-assembling lumazine synthase (LuS) scaffold displaying clade 1a, 1b and 2 RBDs led to a dominant response towards an epitope that overlapped with that of S2H97 [52]. It is thought that multivalent presentation of mixed nanoparticles preferentially expands B cells with cross-reactive BCRs. This approach appears a promising strategy for eliciting nAbs to these conserved but minimally exposed RBD site V and class 4 epitopes.

In summary, the novel V1WT_06, V1WT_41, VA14_26 mAbs are candidates for development of antibody-based therapeutics against newly emerging sarbecoviruses and can provide templates for rational development of pan-sarbecovirus vaccines.

## Acknowledgments

This work was funded by MRC project grant ([MR/X009041/1] to KJD), MRC Genotype-to-Phenotype UK National Virology Consortium ([MR/W005611/1] and [MR/Y004205/1] to KJD and DB), and Wellcome funded consortium - Genotype-to-Phenotype Global ([226141/Z/22/Z] to KJD and DB) and the Francis Crick Institute (to PC), which receives its core funding from Cancer Research UK (CC2058), the UK Medical Research Council (CC2058), and the Wellcome Trust (CC2058). Fondation Dormeur, Vaduz for funding equipment to KJD. KJD was supported by the Medical Research Foundation Emerging Leaders Prize 2021. AU was supported by a PhD Studentship jointly funded by King’s college London and the Francis Crick Institute. ALH was supported by funding from the Bill and Melinda Gates Foundation (Grants OPP1192002, OPP1215550, and INV-035610). DB is supported by a BBSRC Institute Strategic Program Grant (BBS/E/PI/230001A, BBS/E/PI/230002A and BBS/E/PI/230002B) to The Pirbright Institute.

NK-92 cell line was provided by Prof Richard Stanton, University of Cardiff. We gratefully acknowledge all data contributors, i.e., the Authors and their Originating laboratories responsible for obtaining the specimens, and their Submitting laboratories for generating the genetic sequence and metadata and sharing via the GISAID Initiative, on which this research is based.

## Author contributions

Conceptualisation: KJD, PC, JS, AU; Formal analysis: AU, JS, KD, PC, RJO, ALH; Funding acquisition: KJD, PC; Investigation: AU, JS, YA, JN, NT, ALH, JCHT, AN, RJO; Methodology: AU, JS, JCHT, YA, AN, RJO; Project administration: KD, PC; Writing – original draft: KJD, PC, AU, JS; Writing – review and editing: all authors.

## Materials and methods

### Monoclonal antibody isolation and expression

VA14_26 [22], and V1WT_06 and V1WT_41 [8] were isolated previously. IgG1 antibody heavy and light plasmids were co-transfected at a 1:1 ratio into HEK-293F cells (Thermofisher) using PEI Max (1 mg/mL, Polysciences, Inc.) at a 3:1 ratio (PEI Max:DNA). Antibody supernatants were harvested five days following transfection, filtered and purified using protein G affinity chromatography following the manufacturer’s protocol (GE Healthcare).

### RBD expression and purification

Recombinant RBD of SARS-CoV-2, WIV-1, R4231, RacCS203, Rc-o319, RhGB07 for ELISA were expressed and purified as previously described [20]. Briefly, RBDs were cloned into the pOPIN-BAP-His expression vector in Expi293F cells, transfected using PEI40K (Polysciences) with additives (300mM valproic acid, 500mM sodium propionate, 2.5M glucose. Supernatants were harvested 4 days post-transfection and presence of secreted protein was determined by Coomassie and by blotting for anti-His. Supernatants were purified using HisTrapHP columns (Cytivia), desalted using Zeba Spin Desalting Columns (ThermoFisher Scientific) and concentrated using Amicon Ultra 15 columns (Merck Millipore), before quantification by Nanodrop and a Pierce BCA Protein Assay (ThermoFisher Scientific).

### WIV-1 Spike cloning, expression and purification

For cryo-EM, Trimeric WIV-1 spike ectodomain (corresponding to residues 1-1191, GenBank AGZ48831.1) with amino acid substitutions for stabilising the pre-fusion conformation (K969P, V970P) and K668S followed by carboxy-terminal foldon and hexahistidine tag [25] was expressed in HEK-293F cells and purified as previously described [53].

### Fab cloning and expression

The variable heavy and light chain domains were cloned into the pHLsec vector [54] containing the CH1 or CK1 domains, respectively. Antibody heavy and light plasmids were co-transfected at a 1:1 ratio into HEK-293F cells (Thermofisher) using PEI Max (1 mg/mL, Polysciences, Inc.) at a 3:1 ratio (PEI Max:DNA). Antibody supernatants were harvested five days following transfection, filtered and purified using immobilised metal affinity chromatography and size-exclusion chromatography.

### ELISA binding

96-well plates (Corning, 3690) were coated with RBD antigen at 3 μg/mL overnight at 4°C. The plates were washed (5 times with PBS/0.05% Tween-20, PBS-T), blocked with blocking buffer (5% skimmed milk in PBS-T) for 1 h at room temperature. Serial dilutions of IgG or Fab in blocking buffer were added and incubated for 2 hr at room temperature. Plates were washed (5 times with PBS-T) and secondary antibody was added and incubated for 1 hr at room temperature. IgG was detected using Goat-anti-human-Fab-AP (alkaline phosphatase) (1:1,000) (Jackson: 109-055-006). Plates were washed (5 times with PBS-T) and developed with AP substrate (Sigma) and read at 405 nm.

### Pseudotyped virus production

Pseudotyped HIV-1 virus incorporating each of the SARS-CoV-2 variants (D614G, KP.2, LP.1, KP.3, KP.3.1.1, XEC) and sarbecovirus Spikes (SARS-CoV, WIV-1, RhGB07 and Rc-o319) were generated by first seeding HEK293T/17 cells in 10 cm dishes at a density of 3x10^5^ cells/mL in Dulbecco’s Modified Eagle Medium (DMEM) with 10% fetal bovine serum (FBS), 100 U/mL penicillin and 100 µg/mL streptomycin. Following overnight culture, cells were co-transfected using 90 µg PEI-Max (1 mg/mL, Polysciences) with 15 µg HIV-luciferase plasmid, 10 µg HIV 8.91 gag/pol plasmid, and 5 µg Spike protein plasmid [55]. Transfected cells were incubated for 72 h at 37°C, and virus was harvested, sterile filtered, and stored at −80°C until required.

### Neutralisation assays

Neutralisation assays were conducted as previously described [56–58] using either HeLa cells stably expressing the ACE2 receptor (provided by Dr James Voss, Scripps Research, La Jolla, CA) or HEK 293T cells transiently transfected with either Human (provided by Prof Michael Malim, KCL), Halcyon horseshoe bat (*Rh. alycone*) or Japanese horseshoe bat (*Rh. cornutus*) ACE2. HEK293T cells were transfected with either Human, Halcyon or Japanese horseshoe bat ACE2 using PEI Max (1 mg/mL, Polysciences, Inc.) at 3:1, 6:1 and 2:1 ratio (PEI Max:DNA), respectively. Cells were harvested after 72 hours using EDTA.

Serial dilutions of IgG or Fab were prepared with DMEM-C media (10% Fetal Calf Serum and 1% Penicillin/Streptomycin) and incubated with pseudotyped virus for 1-hour at 37°C in 96-well plates. Next, HeLa-ACE2 cells (12,500 cells/50µL per well) or transfected HEK293T/17 cells (15,000 cells/25µL per well) were added and the plates were left for 72 hours. The amount of infection was assessed in lysed cells with the Bright-Glo luciferase kit (Promega), using a Victor™ X3 multilabel reader (Perkin Elmer). Measurements were performed in duplicate and duplicates used to calculate the ID_50_.

### ACE2 competition measured by flow cytometry

Purified mAbs were mixed with His tagged RBD in a molar ratio of 4:1 on ice for 1 h. HeLa-ACE2 cells or HEK293T/17 cells transiently expressing *Rh. alycone*-ACE2 or *Rh. Cornutus*-ACE2 were washed once with PBS and detached using PBS containing 5mM EDTA. Detached cells were washed and resuspended in FACS buffer. Cells (0.5 million) were added to each mAb-RBD complex and incubated on ice for 1 hour. RBDs were stained concomitantly, using FITC-conjugated Rabbit-anti-HIS antibody (Abcam, Ab1206) at 5μg/ml for a final volume of 100μl. The cells were washed with PBS and resuspended in 1 mL FACS buffer with LIVE/DEAD Fixable Aqua Dead Cell Stain Kit (1 μL, Invitrogen). Cells only and cells with RBD only were used as background and positive controls, respectively. The geometric mean fluorescence of FITC was measured from the gate of singlet cells as previously described [59].

ACE2 binding inhibition was calculated with this equation:

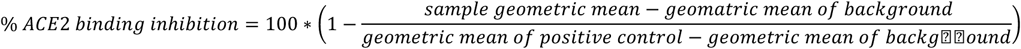

### S1 shedding assay

HEK293T/17 cells were plated in a 6-well plate (2x10^6^ cells/well). Cells were transfected with 1 μg of plasmid encoding full-length SARS-CoV-2 Spike using PEI Max 40K (3 μL at 1 mg/mL) and incubated for 48h at 37°C. Post incubation cells were resuspended in PBS and plated in U-bottom 96-well plates (1x10^5^ cells/well). Monoclonal antibodies were diluted in FACS buffer (1x PBS, 2% FBS, 1 mM EDTA) to 25 μg/ml and incubated with cells on ice for 5, 10, 20, 40 or 80 mins. The plates were washed twice in FACS buffer and stained with 50 μl/well of 1:200 dilution of PE-conjugated mouse anti-human IgG Fc antibody (BioLegend) on ice in dark for 1h. After another two washes, stained cells were analyzed using flow cytometry, and the binding data were generated by calculating the percent (%) PE-positive cells using FlowJo 10 software. Mean fluorescence intensity (MFI) for each sample was determined at each timepoint and each sample was normalised to the MFI at the 5 min timepoint (MFI/MFI at 5 min × 100).

### ADCC assay

NK-92 cells expressing human CD16 were maintained in Minimum Essential Medium-α (MEM-α) supplemented with L-glutamine, nucleosides, 12.5% Foetal Calf Serum, 12.5% Horse Serum, 20mM HEPES, 0.2mM Myo-inositol, 0.02mM Folic Acid, 0.1mM 2-mercaptoethanol and 50 IU/ml Interleukin (IL)-2.

High-binding ELISA plates were coated with recombinant WT Spike protein at 3μg/mL in PBS for 2h at 37 °C. Wells were washed with PBS and then blocked with 5% bovine serum albumin in PBS overnight at 4°C. The plates were washed with PBS, prior to addition of monoclonal antibodies diluted in fully supplemented MEM-α and incubated at room temperature for 1h. NK-92 CD16 cells were added (1.0×10^^5^ cells/well) in media supplemented with 100 IU/ml IL-2, Golgistop protein transport inhibitor (BD Biosciences #554724) and PerCP/Cyanine5.5 anti-human CD107a antibody (Biolegend Clone H4A3). Plates were incubated at 37°C, 5% CO_2_ for 6h. NK-92 CD16 cells were transferred to 75mm Polystyrene Tubes, washed with PBS and fixed by addition of 4% paraformaldehyde. Level of CD107a surface expression was measured by flow cytometry using a BD FACSCanto II.

### Cryo-EM data collection, image processing, and structure refinement

WIV-1 Spike ectodomain (3.5 µl, 1.3 mg/mL), supplemented with V1WT_41, VA14_26, or V1WT_06 Fab (0.2 mg/mL, corresponding to a 1.3-fold molar excess to Spike) in 150 mM NaCl, 0.1% n-octyl glucoside, and 20 mM Tris-HCl, pH 8.0, was applied to 300-mesh 1.2/1.3 C-flat holey carbon grids (Electron Microscopy Sciences) for 1 min under 95% humidity at 20°C and blotted for 3-6 s, and the samples were vitrified by plunge-freezing in liquid ethane/propane using a Vitrobot Mark IV (Thermo Fisher Scientific). Cryo-EM data were acquired across on Titan Krios cryo-electron microscopes with Falcon 4i direct electron detectors (Thermo Fisher Scientific; V1WT_41 and VA14_26). For imaging of the V1WT_06 Fab-Spike complex, the microscope was additionally equipped with a Selectris energy filter with an energy slit set to 10 eV. Micrograph movie stacks (4,500 for V1WT_41 and VA14_26 and 7,058 for V1WT_06 FAB-Spike complex) were recorded in the dose-fractionation mode, at a calibrated magnification corresponding to 1.08 Å (V1WT_41 and VA14_26) or 0.95 Å (V1WT_06) per physical pixel and a total sample exposure dose of 32.2 e/Å^2^ (V1WT_41 and VA14_26) or 41.0 e/Å^2^ (V1WT_06). A defocus range of -1.5 µm to -3.3 µm was used for data collection (**Supplemental Table 1**). 1,674 EER frames, recorded per micrograph movie, were subsequently processed in 31 fractions.

The micrograph stacks were aligned and summed, with dose weighting applied, as implemented in MotionCor2 [60]. Contrast transfer function (CTF) parameters were estimated from image sums using GCTF v1.18 [61]. Initial particle picking was done using crYOLO with general model (gmodel_phosnet_202005_N63_c17.h5) [62]. The resulting 730,359 (VA14_26), 576,253 (V1WT_41), and 1,011,931 (V1WT_06) particles were extracted in RELION-4.0 [63] binned 4-fold to a pixel size of 4.32 Å (V1WT_41 and VA14_26) or 3.8 Å (V1WT_06). These were subjected to four rounds of reference-free 2D classification in cryoSPARC v4.6.2 [64]. Initially, particles belonging to well-defined 2D classes corresponding for the Spike trimer were selected and subjected to ab-initio reconstruction followed by heterogeneous and non-uniform refinement in cryoSPARC (**Supplemental Figure 2**). Neither the 2D nor 3D class averages of the trimeric spike revealed features attributable to a bound FAB molecule (**Supplemental Figure 2**). By contrast, FAB-like features were readily identifiable in 2D class averages corresponding to dissociated Spike (**Supplemental Figure 2**).

Particles contributing to 2D classes of the dissociated Spike were subjected to ab-initio reconstruction followed by heterogeneous refinement in cryoSPARC with 4 classes. Those belonging to the most populated well-resolved 3D classes were used to train a model for picking the entire datasets using Topaz [65]. Newly identified particles extracted binned 4-fold were subjected to three rounds of 2D classification and those contributing to well-defined 2D averages were used for ab-initio reconstruction and, subsequently, heterogenous refinement. The remaining particles were re-extracted with 2-fold binning and subjected to ab-initio reconstruction (5 classes), the obtained reconstructions revealed well-defined 3D reconstructions, readily interpretable as S1 subunit associated with Fab moieties (**Supplemental Figure 2**). Following 3 iterations of heterogeneous refinement in cryoSPARC, 142,232 (VA14_26-S1 complex), 256,418 (V1WT_41-S1 complex) and 135,592 (V1WT_06-S1 complex) particles were re-extracted without binning. These particles were then subjected to ab-initio reconstruction with 5 classes followed by heterogenous refinement in cryoSPARC. Particles from the most populated classes were subjected to Bayesian polishing in Relion-5.0 [66]. Next, ab-initio reconstruction was performed with the polished particles with class similarity set to 0 and number of classes set to 3 (**Supplemental Figure 2**). Subsequently, heterogeneous refinement was conducted using the volume data derived from the ab-initio reconstructions. Local CTF refinement allowed to improve quality of VA14_26-S1 and V1WT_41-S1 maps. The final reconstructions were obtained using non-uniform refinement in cryoSPARC (**Supplemental Table 1, Supplemental Figure 2**). The resolution metrics reported here is according to the gold-standard Fourier shell correlation (FSC) 0.143 criterion [67]. Local resolution was estimated in cryoSPARC (**Supplemental Figure 7**). Illustrations were generated using maps B-factor sharpened maps, generated using cryoSPARC (**Supplemental Figure 5B, 5C and 5E**).

The atomistic model for monomeric S1 protein were derived from PDB entry 8TC0. Antibodies were modelled for their translated amino acid sequence using AlphaFold 3 server (https://alphafoldserver.com/welcome) [68]. Models were docked into cryo-EM maps using UCSF Chimera [69]. Guided by the cryo-EM maps, the models were fitted interactively in Coot-0.9.8.95 [70]. The model was refined using CryoSPARC maps filtered according to local resolution metrics in phenix.real_space_refine (version dev-4213) [71, 72]. The quality of the final model was assessed using MolProbity [73]. The final cryo-EM reconstruction and refined model have been deposited with the EMDB and the PDB under accession codes 9S6Y, 9S6Q and 9S67.

### Analysis of Spike mutations in circulating SARS-CoV-2

The complete SARS-CoV-2 metadata file was downloaded from the GISAID [74] EpiCoV database (https://gisaid.org) on 30/04/2025; details on these sequences is available at https://doi.org/10.55876/gis8.250704no. The file contains an entry for each SARS-CoV-2 sequence including sample information (host, collection date, geographical location, etc) as well as SARS-CoV-2 genome sequence information (length, pangolin lineage, non-synonymous mutations in each ORF, etc). A python script was written to parse this file and extract and count all non-synonymous mutations observed across all Spike amino acid positions; sequences that were not human, were less than 20,000 nucleotides, had an unassigned pangolin lineage or were labelled as “low coverage” were excluded, leaving 15.6 million sequences.

### Surface Plasmon Resonance

Kinetics and epitope binning assays were performed on the Carterra LSA^XT^ using HC30M chips as previously described [20]. Briefly, chips were pre-conditioned using sequential injections of NaOH, NaCl and glycine (pH 2.5). For kinetics, an anti-human Fc lawn was covalently coupled to the HC30M surface using EDC/NHS chemistry. V1WT_06, V1WT_41, and VA14_26 were diluted in PBST (PBS + 0.005 % tween 20) to 1 μg/mL and 0.25 μg/mL and captured to the anti-human-Fc lawn for seven minutes. Single cycle (non-regenerative) kinetics was performed using RBD diluted two-fold in PBST (5 μg/mL to 2.4 ng/mL) allowing a 7-minute association and 14-minute dissociation. Surface regeneration was achieved using glycine (pH 1.9, 10 mM). Kinetic analysis was performed in KineticsTM v1.9.2 to calculate Ka, Kd, and KD. Data was fitted using 2000 iterations with a minimum allowable Kd of 2.5E-05. Average Ka, Kd, and KD were calculated using the values obtained from both coupling concentrations and the two independent couplings on the chip giving n = 4. For epitope binning, human anti-RBD monoclonal antibodies were directly covalently coupled to the HC30M surface. SARS-CoV-2 wild type RBD was injected first followed by the competing antibody. Surface regeneration was achieved as above. Epitope binning analysis was performed in EpitopeTM v1.9.2.

