## Supplemental Figures and tables for "Mechanisms for cross-neutralisation of diverse bat sarbecoviruses"

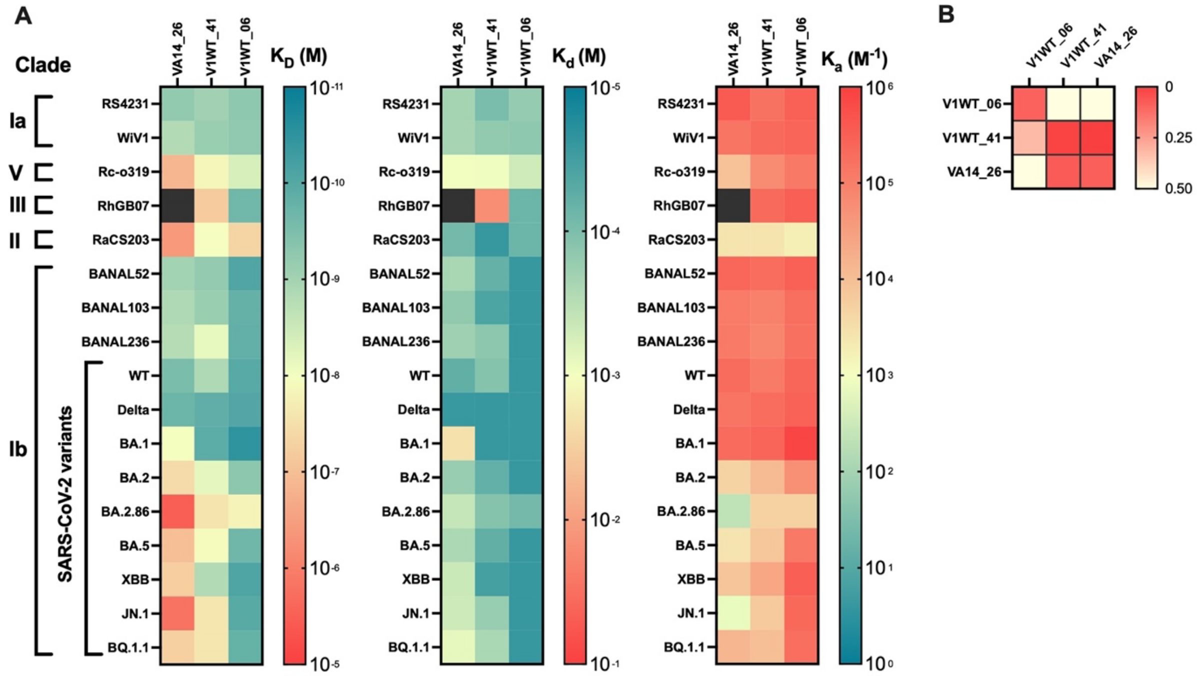


**Supplemental Figure 1: Antibody binding affinity and kinetics and antibody binding competition. A)** Affinity ($K_D$), association rate constant ($k_a$) and dissociation rate constant ($k_d$) of mAb binding kinetics for SARS-CoV-2 RBD were measured by surface plasmon resonance using the Carterra LSA platform. Values are coloured an accordance with the gradients presented. **B)** Heatmap of antibody blocking and sandwiching interactions between VA14_26, V1WT_41 and V1WT_06 measured using SPR against SARS-CoV-2 RBD. Antibodies immobilised as ligands are listed on the left and analytes injected are listed across the top. Scale shows the normalised change in response units (RU) relative to injection of PBS after RBD. RU >0.3 show sandwiching analytes (i.e. no competition) and RU <0.3 shows blocked or displaced analytes (i.e. competition).

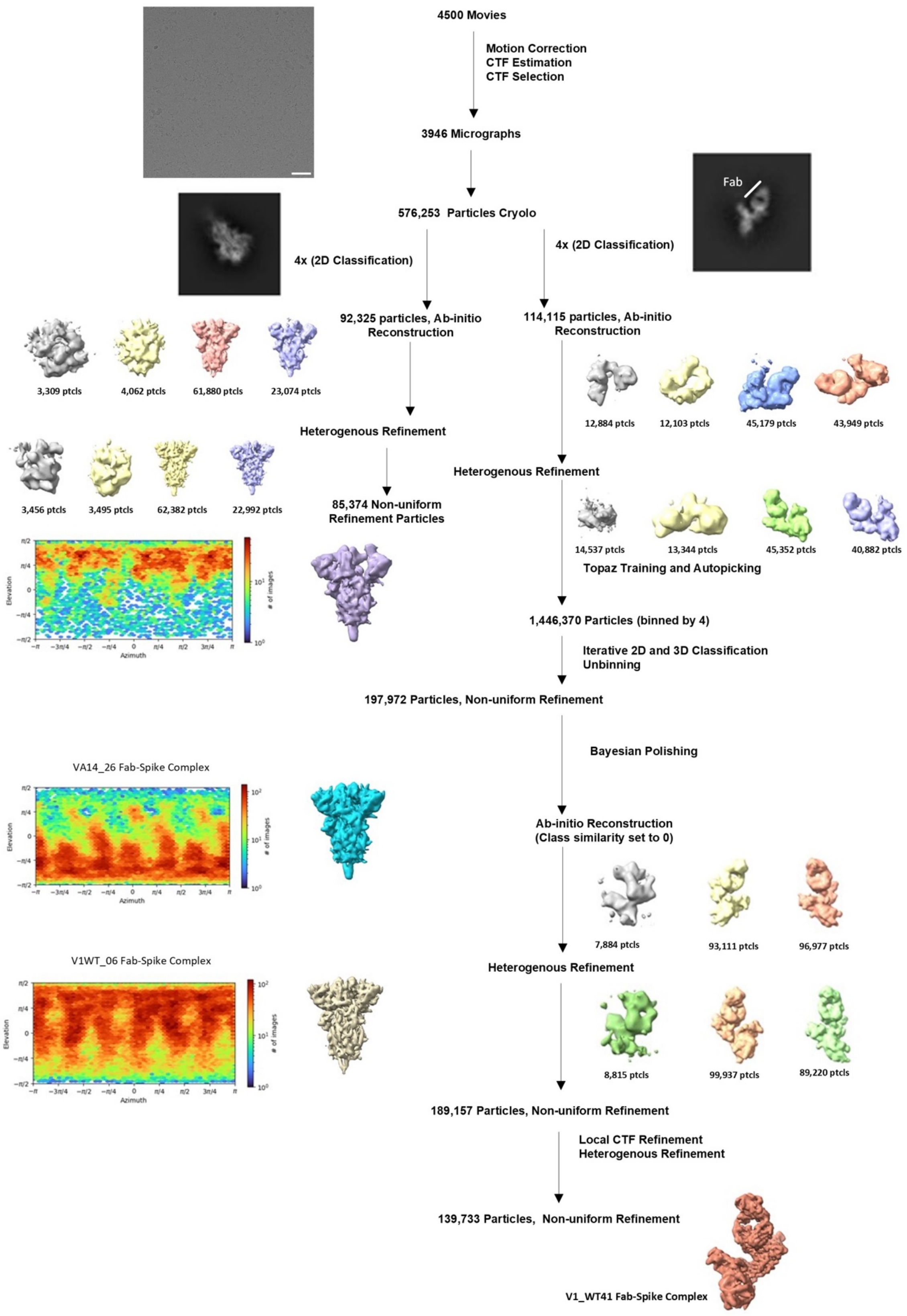


**Supplemental Figure 2: Representative workflow of V1WT_41 Fab-spike complex dataset.** Representative micrograph shown (scale bar at 50 nm). Representative 2D class average corresponding to Spike trimer ectodomain is displayed. Non uniform refinement of the WIV-1 ectodomain trimer reconstruction presented next to viewing direction distribution plot for all three datasets. Notably features corresponding to a bound Fab molecule are absent in 2D and 3D classes. Representative 2D class averages corresponding to dissociated spike protomer bound to V1WT_41 Fab is shown. Abbreviations used: Ptcls – Particles

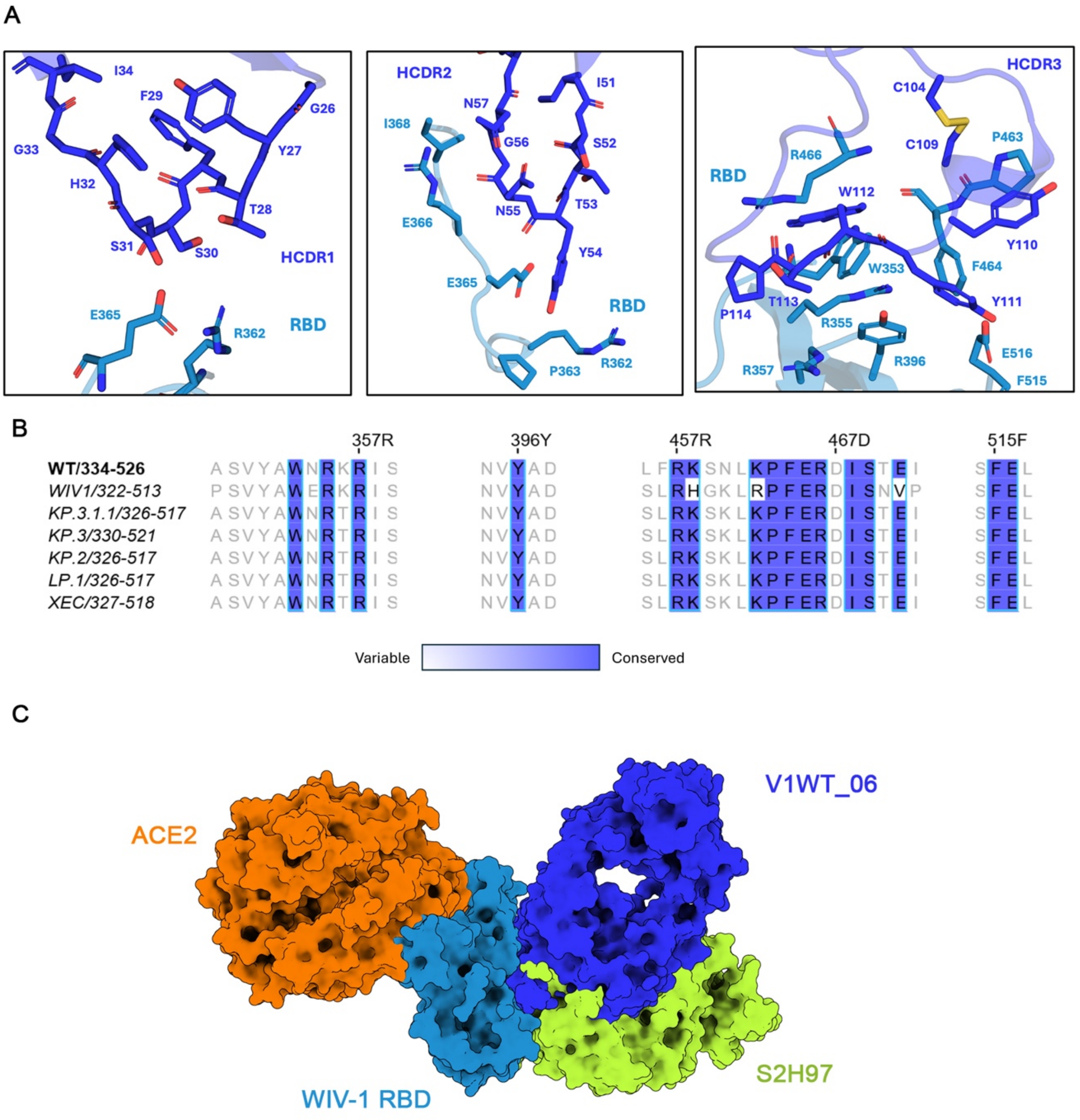


**Supplemental Figure 3. Binding of V1WT_06 to WIV-1 RBD. A)** Closeup view of V1WT_06 HCDRs and the WIV-1 RBD interface. Individual CDRH1, CDRH2 and CDRH3 views have been shown with contact residues on the RBD shown as sticks and renumbered based on alignment to SARS-CoV-2. **B)** Sequence alignment of a panel of SARS-CoV-2 Variant RBDs. Contact residues of V1WT_06 mAb on the WIV-1-RBD are indicated with black outline. The colouring scheme describes the percent identity (PID) between the sequences as implemented in Jalview 2.11.4.0. **C)** Superimposed 3D structure of S2H97 (Lime), another RBD site V neutralising antibody (from PDB: 7M7W) and ACE2 (orange, from PDB: 8WLZ)

shown around WIV-1 RBD – V1WT_06 (dark blue) Fab complex. Angle of approach of V1WT_06 in relation to S2H97 contrasts the RBM proximal, heavy chain driven interaction of V1WT_06 as opposed to RBM distal angle of approach of S2H97 [1].

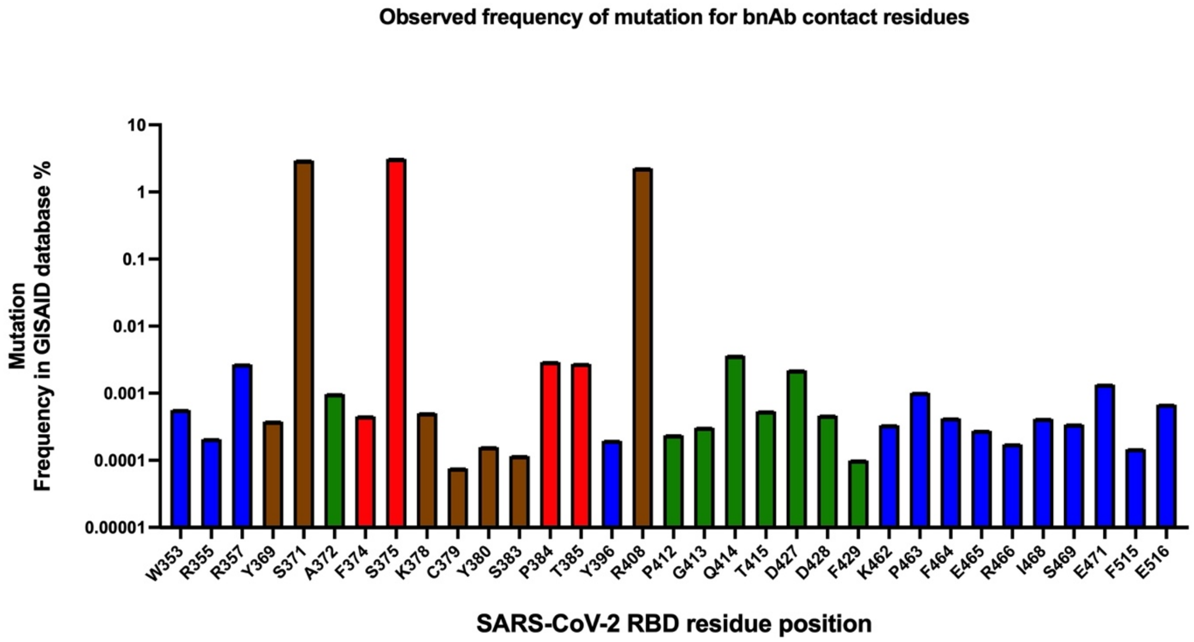


**Supplemental Figure 4. Conservation of mAb contact residues observed in the GISAID data base.** Frequency of mutations at RBD contact residues observed in the GISAID database. Residue position is based on SARS-CoV-2 numbering. Contact residues are coloured according to the mAb; V1WT_06 (blue), VA14_26 (red), V1WT_41 (green) and shared VA14_26/V1WT_41 contact residues (brown).

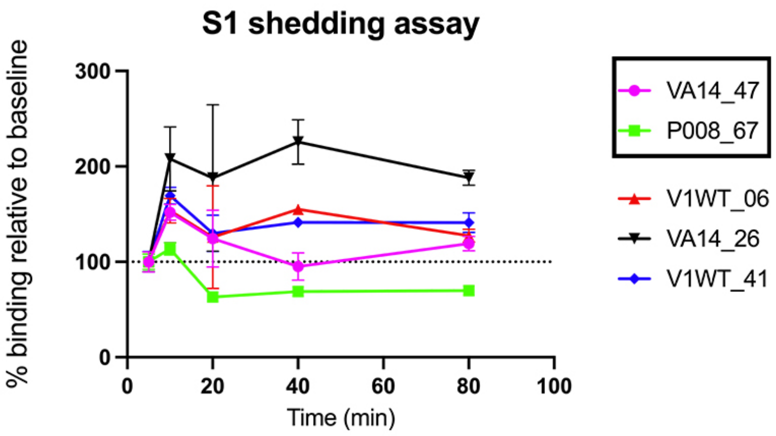


**Supplemental Figure 5: S1 shedding as a mechanism of SARS-CoV-2 neutralisation.** Antibody mediated shedding of S1 from SARS-CoV-2 Spike. HEK 293T cells expressing Wuhan-1 Spike were incubated with mAbs and binding measured by flow cytometry at 5, 10, 20, 40 and 80 mins. mAb P008_67 was used as a positive control and VA14_47 was used as a negative control [2]. The SARS-CoV-2 Spike protein does not contain the furin cleavage site mutation.

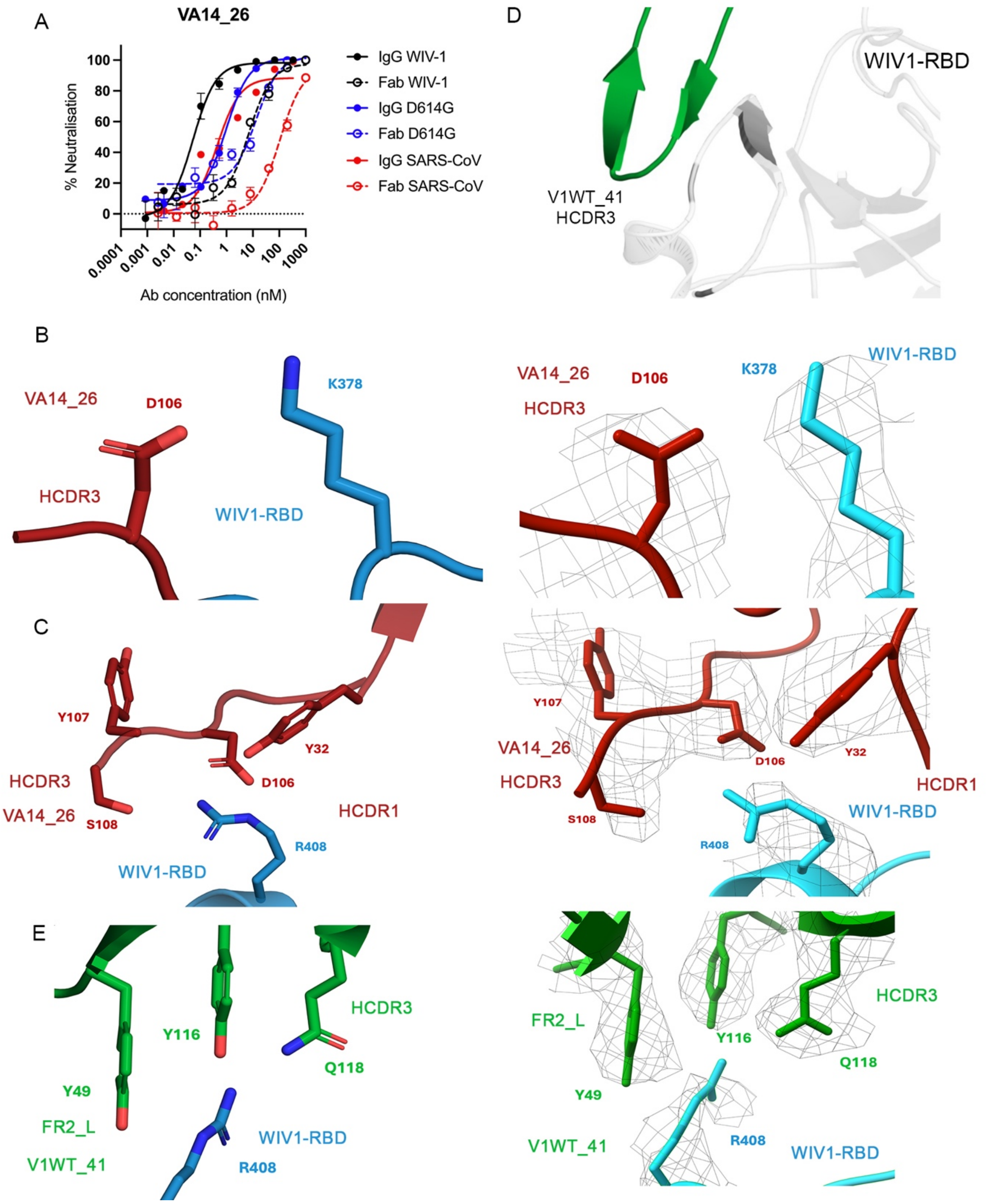


**Supplemental Figure 6. Binding of VA14_26 and V1WT_41 to WIV-1 RBD. A)** Comparison in neutralisation potency of VA14_26 Fab and IgG against SARS-CoV-2 (D614G), SARS-CoV-1 and WIV-1. **B)** Close up view of K378 on WIV-1 RBD and its contact residue on VA14_26. Panel on the right depicts the coulomb density mesh around the residues at SD level 9, as implemented in ChimeraX_1.7.1. **C)** Close up view of R408 on WIV-1 RBD and its contact residues on VA14_26. Panel on the right depicts the coulomb density mesh around the residues at SD level 10.25, as implemented in ChimeraX_1.7.1. **D)** Close up view of YYDRSG motif in V1WT_41 HCDR3 (green) in contact with WIV-1 RBD (grey), shown in cartoon representation. Regions outside the contact residues are turned transparent. Local structure with right-handed twist of beta hairpin is visible. **E)** Close up view of R408 on WIV-1 RBD and its contact residues on V1WT_41. Panel on the right depicts the coulomb density mesh around the residues at SD level 12.5, as implemented in ChimeraX_1.7.1.

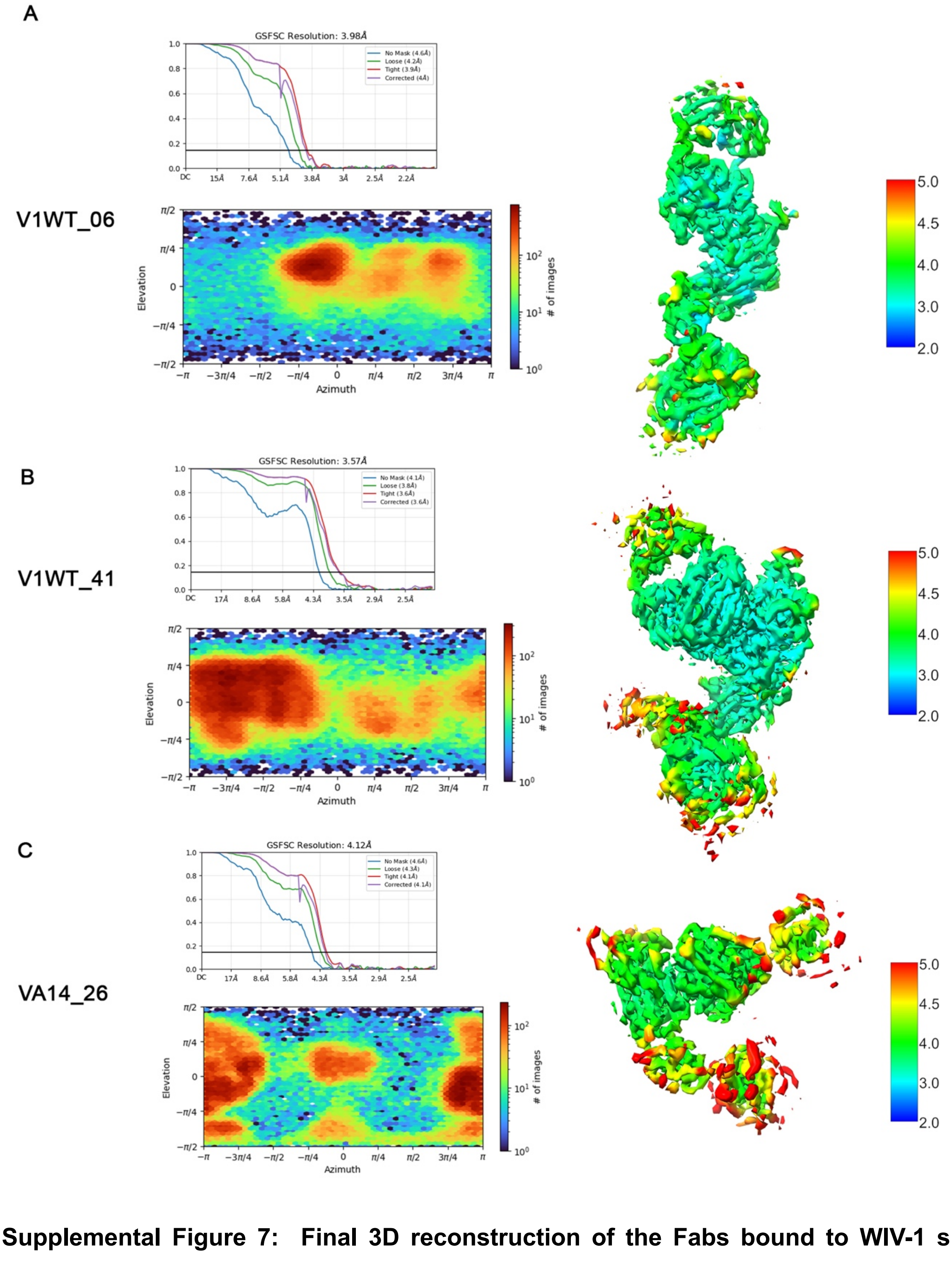


**Supplemental Figure 7: Final 3D reconstruction of the Fabs bound to WIV-1 spike protomer. A)** Panel displaying Half-map Fourier shell correlation curve (FSC curve) and viewing direction distribution plot (cryoSparc) for final reconstruction of V1WT_06 Fab-Spike protomer complex. The reconstruction is coloured by estimated local resolution (cryoSparc), colour scheme provided on the right. **B)** Panel displaying Half-map Fourier shell correlation curve (FSC curve) and viewing direction distribution plot (cryoSparc) for final reconstruction of V1WT_41 Fab-Spike protomer complex. The reconstruction is coloured by estimated local resolution (cryoSparc), colour scheme provided on the right. **C)** Panel displaying Half-map Fourier shell correlation curve (FSC curve) and viewing direction distribution plot (cryoSparc) for final reconstruction of VA14_26 Fab-Spike protomer complex. The reconstruction is coloured by estimated local resolution (cryoSparc), colour scheme provided on the right.

**Supplemental Table 1:** Cryo-EM data collection, refinement and validation statistics

| | **VA14_26–spike** complex | **V1WT_41– spike** complex | **V1WT_06–spike** complex |
|---|---|---|---|
| **Data collection and processing** | | | |
| Microscope/ detector | Krios1/Falcon 4i | Krios1/Falcon 4i | Krios 2/Falcon 4i |
| Grid type | 300-mesh C-flat holey carbon grids (CF 1.2/1.3 ) | 300-mesh C-flat holey carbon grids (CF 1.2/1.3 ) | 300-mesh C-flat holey carbon grids (CF 1.2/1.3 ) |
| Voltage (kV) | 300 | 300 | 300 |
| Total exposure (e/Å) | 32.2 | 32.2 | 42 |
| Energy filter slit (eV) | - | - | 10 |
| Movies acquired | 4,500 | 4,500 | 7,058 |
| Defocus range (µm) | -1.5 to -3 | -1.5 to -3 | - 1.5 to -3.3 |
| Exposure time (s) | 5.44 | 5.44 | 5.44 |
| Pixel size (Å) | 1.08 | 1.08 | 0.95 |
| Number of EER frames | 1,674 | 1,674 | 1,674 |
| Calibrated Magnification | 75,000 | 75,000 | 130,000 |
| Dose rate ( e/Å$^2$/s) | 5.92 | 5.92 | 7.53 |
| Total dose/Frame (e/Å$^2$) | 32.2 | 32.2 | 41.0 |
| Particles picked (Topaz autopick) | 1,460,644 | 1,446,370 | 3,514,633 |
| Particles used in final reconstruction | 86,961 | 139,733 | 101,825 |
| Map resolution (Å) | 4.12 | 3.57 | 3.98 |
| FSC threshold | 0.143 | 0.143 | 0.143 |
| **Refinement** | | | |
| CCbox | 0.71 | 0.68 | 0.79 |
| CCmask | 0.70 | 0.78 | 0.74 |
| Bond Length (Å) | 0.003 | 0.003 | 0.003 |
| Bond Angle (°) | 0.519 | 0.512 | 0.509 |
| Ramachandran favoured (%) | 95.45 | 94.50 | 96.13 |
| Ramachandran outliers (%) | 0 | 0 | 0 |
| MolProbity clash score | 6.51 | 2.60 | 5.23 |
| MolProbity overall score | 1.93 | 1.59 | 1.74 |
